# Bcl11b is a Newly Identified Regulator of Vascular Smooth Muscle Phenotype and Arterial Stiffness

**DOI:** 10.1101/193110

**Authors:** Jeff Arni C. Valisno, Pavania Elavalakanar, Christopher Nicholson, Kuldeep Singh, Dorina Avram, Richard A. Cohen, Gary F. Mitchell, Kathleen G. Morgan, Francesca Seta

**Affiliations:** Vascular Biology Section, Department of Medicine, Boston University School of Medicine, Boston, MA, USA; Department of Health Sciences, Sargent College, Boston University, MA, USA; Department of Medicine, University of Florida, FL, USA; Cardiovascular Engineering, Norwood, MA, USA

**Keywords:** vascular smooth muscle cells, arterial stiffness, vascular function, aortic aneurysm

## Abstract

B-cell leukemia 11b (Bcl11b) is a zinc-finger transcription factor known as master regulator of T lymphocytes and neuronal development during embryogenesis. Bcl11b-interacting protein COUP-TFII is required for atrial development and vasculogenesis, however a role of Bcl11b in the adult cardiovascular system is unknown. A genome-wide association study (GWAS) recently showed that a gene desert region downstream of *BCL11B* and known to function as *BCL11B* enhancer harbors single nucleotide polymorphisms (SNPs) associated with increased arterial stiffness. Based on these human findings, we sought to examine relations between Bcl11b and arterial function using mice with Bcl11b deletion. We report for the first time that Bcl11b is expressed in vascular smooth muscle (VSM) and transcriptionally regulates the expression of VSM contractile proteins smooth muscle myosin and smooth muscle α-actin. Lack of Bcl11b in VSM-specific Bcl11b null mice (BSMKO) resulted in increased expression of Ca^++^-calmodulin-dependent serine/threonine phosphatase calcineurin in BSMKO VSM cells, cultured in serum-free condition, and in BSMKO aortas, which showed an inverse correlation with levels of phosphorylated VASP^S239^, a regulator of cytoskeletal actin rearrangements. Moreover, decreased pVASP^S239^ in BSMKO aortas was associated with increased actin polymerization (F/G actin ratio). Functionally, aortic force, stress and wall tension, measured ex vivo in organ baths, were increased in BSMKO aortas and BSMKO mice had increased pulse wave velocity, the *in vivo* index of arterial stiffness, compared to WT littermates. Despite having no effect on baseline blood pressure or angiotensin II-induced hypertension, Bcl11b deletion in VSM increased the incidence of aortic aneurysms in BSMKO mice. Aneurysmal aortas from angII-treated BSMKO mice had increased number of apoptotic VSM cells. In conclusion, we identified VSM Bcl11b as a novel and crucial regulator of VSM cell phenotype and vascular structural and functional integrity.

## INTRODUCTION

A recent genome-wide association study (GWAS) of 9 discovery cohorts (20,634 participants; average age of cohort 34-75 years old) and 2 replication cohorts (N=5,306) identified single nucleotide polymorphisms (SNPs) in a gene desert locus on chromosome 14 with highly significant association (p < 5.6 x 10^-11^ for the top SNP, rs1381289T>C) ^1^ with pulse wave velocity (PWV), the gold standard clinical measure of aortic wall stiffness ^2^. Notably, the presence of each rs1381289 allele resulted in an estimated hazard ratio (HR) = 1.05 for a first major coronary artery disease event and HR = 1.10 for incident heart failure, suggesting that increased arterial stiffness associated with SNP variants in this locus may be causally linked to an increased risk of subsequently developing a major cardiovascular disease event.

The “aortic stiffness” locus spans 2 Mb between coding genes B-cell leukemia 11b (*BCL11B*) and vaccinia-related kinase 1 (*VRK1*). A detailed analysis indicated that the region contains an ∼1.9 kb sequence, located ∼850 kb downstream (3’) of *BCL11B*, known to function as an enhancer for *BCL11B* ^3^ but not *VRK1*. Enhancers are important DNA regulatory elements that activate the expression of target genes, independently of distance or orientation ^4^. In addition, the long non-coding RNA (lncRNA) DB129663, of unknown biological function, partially overlaps with the *BCL11B* enhancer (Figure 1). In the present study, we sought to elucidate the mechanistic basis of the association of the chromosome 14 locus with aortic stiffness and a possible cause-effect relationship between Bcl11b and vascular function.

**Figure 1.**
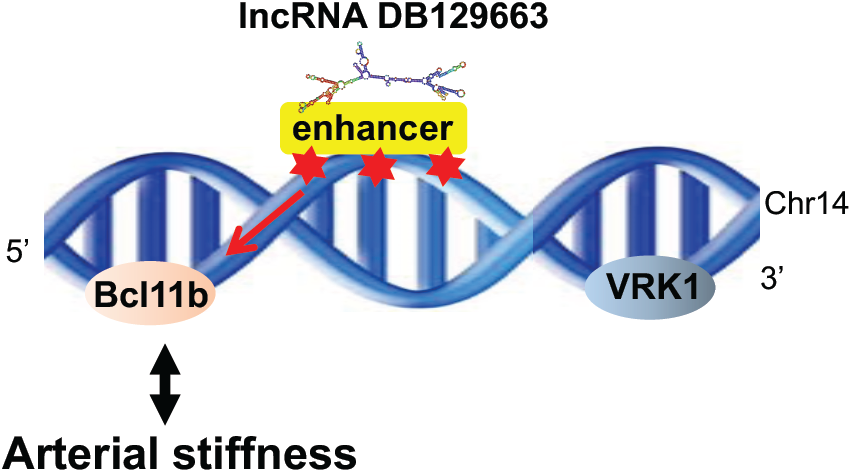
Genetic locus of genome-wide association with arterial stiffness. A gene desert region on chromosome 14, between genes *BCL11B* and *VRK1*, contains a *BCL11B* enhancer (*in yellow*) and an overlapping lncRNA DB129663, where SNP variants (*red stars*) associated with increased arterial stiffness are present.

Bcl11b, also known as COUP-TF interacting protein-2 (CTIP2), is a zinc finger transcription factor ^5^ best known as a regulator of T cell lineage commitment during hematopoiesis ^6–8^ and neuronal development during embryogenesis ^9,10^. However, a role of Bcl11b in the vasculature has never been described. Here we report for the first time that Bcl11b is expressed in human and murine aortas and it is a crucial determinant of vascular smooth muscle (VSM) contractile phenotype, aortic wall stiffness and aortic structural integrity. Bcl11b transcriptionally regulates VSM-specific gene expression, including contractile proteins smooth muscle myosin and smooth muscle α-actin and the Ca^++^-calmodulin-dependent serine/threonine phosphatase calcineurin. Bcl11b deletion in VSM cells (BSMKO mice) resulted in increased actin polymerization (F/G actin ratio), increased baseline aortic tone and increased PWV compared to WT littermates. No significant blood pressure differences were noted between WT and BSMKO mice at baseline or upon ang II administration. However, 2-week ang II treatment induced aortic aneurysms in BSMKO mice with a ∼ 70% incidence. Taken together, our study identifies VSM Bcl11b as a novel and crucial regulator of VSM functional and structural integrity. Bcl11b represents a promising candidate for the translational development of therapies against arterial stiffness, a major risk factor for cardiovascular disease, and aortic aneurysms.

## METHODS

### Mice with VSM-specific Bcl11b deletion

Animal studies were approved by the Boston University Medical Center Institutional Animal Care and Use Committee (IACUC) in accordance with federal regulations and guidelines on the humane use of experimental animals.

Floxed Bcl11b mice (Bcl11b^flox/flox^) in which Bcl11b’s exon 4, crucial for its zinc finger DNA binding activity, is flanked by loxP sites ^8^, were inter-crossed with transgenic mice in which Cre-recombinase is linked to an estrogen receptor construct for tamoxifen-inducible removal of Bcl11b (ER-Cre-Bcl11b^flox/flox^) and WT littermate controls. In order to study the functional role of Bcl11b in VSM cells, we bred Bcl11b^flox/flox^ mice with transgenic mice in which Cre-recombinase expression is driven by smooth muscle myosin heavy chain 11 (*SMMHC* or *MYH11*) promoter linked to an estrogen receptor construct (SMMHC^CreERT2^, stock number 019079, The Jackson Laboratory), which allows for tamoxifen-inducible removal of Bcl11b in VSM. Eight-week old ER-Cre-Bcl11b^flox/flox^ or SMMHC^CreERT2^/Bcl11b^flox/flox^ mice received tamoxifen (200μl, 2mg/d, 5 days) or vehicle (200μl, sunflower oil, 5 days) to obtain global Bcl11b null mice (BKO) and VSM-specific Bcl11b-deficient mice (BSMKO) and WT littermate controls, respectively. WT, BKO and BSMKO mice entered study protocols no earlier than 2 weeks after the last tamoxifen or vehicle administration. In addition, to corroborate Bcl11b vascular effects in a second animal model, we inter-crossed Bcl11b^flox/flox^ with transgenic mice in which Cre-recombinase expression is driven by endogenous smooth muscle 22 α protein promoter (SM22α, transgelin; stock number 006878, The Jackson Laboratory) to generate mice with constitutive Bcl11b deletion in VSM (SM22α-Cre^+^/Bcl11b^flox/flox^ and SM22α-Cre^-^/Bcl11b^flox/flox^ littermate controls), as we previously described^11,12^. Mice were backcrossed into C57Bl/6J genetic background for at least 10 generations.

### *In vivo* arterial stiffness measurements

Pulse wave velocity (PWV), the gold standard *in vivo* measure of aortic wall stiffness, was measured in (1) 10-months old WT (n=4) and global Bcl11b-deficient mice (n=4), and (2) WT (n=14) and VSM-specific Bcl11b-null mice (n=17; 9 BSMKO, 2 months after tamoxifen administration, and 8 SM22αBKO, at 2 months of age), by methods that we described previously ^11–15^. Briefly, mice were kept recumbent and lightly anesthetized with 1-2% isofluorane while on a heated platform that maintained body temperature during the procedure. Pressure or flow pressure waves and simultaneous electrocardiogram (ECG) were obtained from two locations along the aorta, using a dual-pressure high-fidelity pressure transducer (Mikro-tip catheter transducers, SPR-1000, Millar Instruments, Houston, TX, USA) interfaced with an acquisition and analysis workstation (NIHem, Cardiovascular Engineering, Norwood, MA, USA) or high-resolution Doppler ultrasound (Vevo2100, Fujifilm Visualsonics, Toronto, Canada). Arrival times of the pressure or flow waves at proximal and distal locations were measured by the foot-to-foot method using the R-wave of the ECG as fiducial point, on at least 5 cardiac cycles for each mouse. PWV was calculated as a ratio of the distance and the difference in arrival times of pressure or flow waves at the two locations (m/s). The investigator performing PWV measurements and post-acquisition calculations was blinded to mice genotype. We previously optimized PWV measurements in mice so that mean arterial pressure (MAP = 95 ± 5 mmHg) and heart rate (HR = 450 ± 50 bpm) are maintained constant and at comparable levels between mice^11–15^ because PWV is dependent on both distending pressure (MAP) and heart rate ^16^. However,in initial experiments with global Bcl11b-deficient mice, we intentionally increased MAP by jugular vein infusions of phenylephrine (100 μl, 10 μg/ml) in order to measure PWV over a range of MAP 80-140 mmHg to exclude any possible cofounding effects of MAP on PWV.

### Isometric tension measurements

of aortic rings from WT (n=5) and BSMKO (n=7) mice were performed as we previously described ^14,17,18^. Briefly, aortic rings (∼ 4 mm axial length) were mounted on triangular wires attached to a force transducer, which recorded vessel wall tension. Each ring was stretched to optimal length L_O_ (1.8 × slack length) ^18^ and left to equilibrate for 30 minutes in organ baths containing oxygenated (95% O_2_-5% CO_2_) physiological salt solution (PSS Krebs solution – in mM: 120 NaCl, 5.9 KCl, 1.2 NaH_2_PO_4_, 25 NaHCO_3_, 11.5 dextrose, 1 CaCl_2_, and 1.4 MgCl_2_; pH = 7.4). VSM viability was confirmed by depolarization through the addition of PSS, containing 51 mM KCl instead of NaCl, followed by return to PSS for 15 minutes. Force measurements were acquired for 15 minutes using LabChart software (ADInstruments, Sidney, Australia). Prior to mounting the rings on the wires, aortic ring length and diameter were measured under a light microscope. Wall thickness was measured as the difference between the external and internal elastic laminae from digital images acquired with a fluorescence microscope (20x; NIS-Elements software, Nikon Instruments), using the autofluorescence of elastin fibers and after incubating aortic rings in nuclear stain (NucBlue, Life Technologies). Wall tension (N/m) was calculated as the ratio between force and vessel length; stress (kPa) was calculated as the ratio between force and cross-sectional area. Measurements were acquired in a genotype-blinded fashion.

### Blood pressure measurements

Blood pressures (systolic, mean, diastolic, pulse pressures and heart rate) were measured in WT (n=7) and BSMKO (n=8) mice by radiotelemetry, the gold standard measure of blood pressure in experimental animals, using standard methods (transmitter model TA11PA-C10, Data Sciences International, Saint Paul, MN, USA), as we described previously ^12,13^. After full recovery from surgical implantation of telemetry transducers (7 days), baseline blood pressures were obtained for 8 days, with a schedule of 1 recording every 4 minutes generating 360 daily recordings per mouse. Mice were subsequently implanted with an osmotic minipump (Alzet, Durect, Cupertino, CA, USA) to receive continuous infusions of angiotensin II (1 μg/kg/min) for the following 2 weeks. Baseline and post-angiotensin II blood pressure values were calculated by averaging daily data-points for each mouse in a blinded fashion, before performing statistical analysis by genotype or treatment group.

### Isolation and treatments of VSM cells

VSM cells were isolated from aortas of WT and BSMKO mice by enzymatic dissociation with 1 mg/ml elastase and 3 mg/ml collagenase II, as we described previously ^11,12^. Freshly isolated VSM cells were cultured in DMEM, 1 g/L glucose, 1% antibiotic-antimycotic and 10% fetal bovine serum (FBS) until passage 2-3. In different sets of experiments, VSM cells were made quiescent by using DMEM without FBS for 72 hours and before receiving the following treatments: vehicle control, TGFβ (20 ng/ml, overnight) or cyclosphorine A (1 or 10 μM, overnight). At the end of treatment periods, VSM cells were collected in RIPA or Triazol lysis buffer for protein or mRNA analysis, respectively. In replicate experiments for each condition tested, we used VSM cells freshly isolated from different WT and BSMKO mice to ensure not only technical replicates but also biological replication (i.e, cell preparations from different mice).

### Transfection of aortic rings with Blc11b plasmid and with DB129663 silencing RNA

Aortas were gently stripped of the adventitia after a 12-minute treatment with 3 mg/ml collagenase II in 1 ml DMEM at 37°C. The aortic media layer was then cut in 2 mm rings, which were cut open and incubated for 24 hrs at 37°C with 20 μg of full-length Bcl11b plasmid (clone ID MmCD00316071, Harvard Medical School, Boston) mixed with Lipofectamine 3000 transfection reagent in 500 μl of DMEM with no FBS and no antibiotics. The next day, the medium was replaced with fresh DMEM with 10% FBS for 72 hours. Aortic rings were then briefly washed with cold PBS, snap-frozen in liquid N_2_ and kept at -80°C until further processing for protein analysis.

Human embryonic kidney (HEK) cells were grown in triplicate wells of 6-well plates with DMEM containing Glutamax and 10% FBS at ∼ 70% confluency. HEK cells were transfected with control (non-targeting) siRNA or a mixture of 4 custom-designed silencing RNAs (1.6 μM), each targeting exons 1, 2 or 3 of DB129663 (Sigma-Aldrich, St. Louis, MO, USA), with Lipofectamine RNAiMax in DMEM (no FBS and no antibiotics). After 6 hours, culture medium was replaced with full medium for further 48 hours. Cells were collected in Trizol and kept -80°C until further processing for RNA extraction and qRT-PCR.

### Bcl11b, DB129663, MYH11, α-SMA, MYOCD, ki-67 mRNA quantitation by qRT-PCR

VSM cells, treated as above, were homogenized in Trizol and kept at -80°C until RNA extraction with a commercially available column-based RNA purification kit (ZymoResearch, Irvine, CA, USA), following manufacturer’s recommendations. RNA quality (18S/28S ribosomal RNA ratio) was analyzed with a Bioanalyzer (Boston University Microarray Core) and samples with a RIN ≥ 7 were deemed of good quality and further processed. Messenger RNA was retrotranscribed into cDNA with High capacity RNA-to-DNA kit (Life Technologies), as per manufacturer’s instructions. In addition, cDNA from aortas of the following experimental groups were processed for qRT-PCR: (1) mice fed normal (ND; n=8) or high fat, high sucrose (HFHS; n=10) diet for 8 months, as we described previously ^11,13^; (2) young (5 months old; n=4) and aged (29-month old; n=4) mice, and (3) 9 human subjects (obtained from Dr. O’Shaughnessy, University of Cambridge, UK). Custom-designed primers for lncRNA DB129663 and the mouse homolog AI060616 (Exilarate LNA^™^, Exiqon, Woburn, MA, USA) were tested for specificity by Sanger sequencing of purified PCR products, which confirmed DB129663 and AI060616 nucleotide sequence (Tufts University Sequencing Core). *BCL11B*, myosin heavy chain 11 (*MYH11*), smooth muscle α-actin (*ACTA2*), myocardin (*MYOCD*), *ki-67* gene expressions were analyzed with validated Taqman assays multiplexed for β-actin, in AB750 instrument (Applied Biosystems, Boston University Analytical Core). Relative mRNA expression was calculated by the ΔΔCt method and expressed as fold ratio versus WT controls, with β-actin serving as endogenous control.

### Western blot on aortas and VSM cells

Aortas and VSM cells were manually homogenized in a glass-glass tissue grinder (Kontes, number 20) in RIPA buffer (100-200 μl, Cell Signaling Technology, CST, Danvers, MA, USA), freshly supplemented with protease inhibitors. Cell or tissue homogenates were sonicated on ice (10 seconds, x3) then cleared of non-soluble material by centrifugation at 13,200 rpm for 15 min at 4°C. Protein concentration was measured by a standard colorimetric method with a Pierce BCA Protein Assay kit (Thermo Scientific, Rockford, IL, USA), as per manufacturer’s instructions.

Thirty μg of VSM cell or aortic proteins were resolved by SDS-PAGE in 4-15% polyacrylamide gels (100V, 60 min) and transferred to PVDF membranes (130 mA, 120 min). Membranes were blocked in 5% non-fat milk in 1% Tween-20 in TBS (TBS-T) for 1 hour and incubated overnight with one of following antibodies at 1:1000 or 1:5000 dilution: Bcl11b (catalog# ab18465, Abcam), MYH11 (catalog # ab53219, Abcam), α-SMA (catalog # ab5694, Abcam), phosphorylated VASP at serine 239 (catalog # 3114, CST), total VASP (catalog # 3132, CST), calcineurin (catalog # 2614, CST), β-actin (catalog # 4970, CST) and GAPDH (catalog # 2118S, CST). Membranes were washed in TBS-T and incubated with an appropriate peroxidase-conjugated anti-rat or anti-rabbit secondary antibody (CST) for 1 hour, washed in TBS-T (8 minutes, x3) and exposed to the chemiluminescent substrate ECL (GE Healthcare, Boston, MA, USA) for 5 min to visualize protein bands. In some cases, membranes were incubated for 30 min at room temperature in Restore Plus Western Blot Stripping Buffer (ThermoFisher Scientific, Waltham, MA, USA) to re-probe proteins with a different antibody. The membrane was then blocked for 1 hr in 5% non-fat milk TBS-T and incubated with the respective primary antibody, following the same procedures to visualize protein bands. Blot images were acquired with ImageQuant LAS4000 (General Electric Healthcare Lifesciences, Pittsburgh, PA, USA), which uses a CCD camera set at -25°C and automatic optimal exposure. Protein band intensities were quantified with Image J (www.imagej.nih.gov) and normalized to β-actin or GAPDH, which were used as loading controls. Band intensities pertaining to the same experimental group were averaged and expressed in arbitrary units (A.U.) relative to WT controls.

### Immunohistochemical stainings of aortic sections

Freshly isolated aortic segments (thoracic proximal, thoracic distal and abdominal aorta) from 3 mice were aligned side-by side in OCT so that each slide contained 3 sections, each representing an anatomical segment of an individual mouse aorta. OCT blocks were snap-frozen in liquid N_2_ and cross-sectioned at a thickness of 10μm. After washing in PBS (5 minutes, x3), aortic sections were incubated for 1 hr in 5% milk/TBT-T, to block non-specific binding, and then overnight with 5% BSA/TBT-T containing anti-Bcl11b, at 1: 200 dilution or with rabbit IgG, which served as a negative control for antibody specificity. The next day, sections were washed in TBT-T (5 minutes, x3), incubated for 1 hr with anti-rabbit Texas red-conjugated IgG, washed again with TBT-T (5 minutes, x3) and mounted with DAPI-containing medium Vectashield Hardset (Vector Laboratories) and glass coverslips. Aortas, lungs, hearts and kidneys from tamoxifen-treated ER-Cre-Bcl11b^flox/flox^-mT knock in mice, which aid the visualization of Bcl11b endogenous localization, were similarly processed and imaged except that antibodies were omitted because Bcl11b was already fluorescently tagged (mTomato). Fluorescent digital images were acquired with a Nikon Epifluorescence Microscope (Eclipse 80i) and NIS-Elements 3.22 software, at 20x and 40x magnification (Boston University Cellular Imaging Core).

### Terminal deoxynucleotidyl transferase (TdT) nick-end labeling (TUNEL) of aortic sections

Aortic cryosections from control and angII-treated WT and BSMKO mice were fixed with 4% paraformaldhehyde for 15 minutes at room temperature and washed twice with PBS for 5 minute each. Aortic sections were then processed for TUNEL staining as per manufacturer’s recommendations (Mebstain, Apoptosis TUNEL Kit Detect, MBL, Nagoya, Japan). Briefly, aortic sections were equilibrated in TdT buffer (proprietary manufacturer’s formulation) and incubated for 1 hr at 37°C with 50 μl mixture of TdT enzyme, TdT buffer and FITC-conjugated dUTP. At the end of the incubation, slides were washed in distilled water (2 minutes, x4), counterstained with propidium iodide to visualize nuclei, and mounted with FluorSafe^™^ Reagent (EMD Millipore, Billerica, MA, USA). FITC-positive nuclei, indicative of incorporated dUTP in fragmented DNA of apoptotic cells, were detected with the Nikon Epifluorescence Microscope and quantified with Image J. A positive control slide was prepared by incubating a WT aortic or liver section with DNase I (5μl, 6U/μl; ZymoResearch) for 30 minutes at room temperature and then processed as above, together with other slides.

#### F/G actin quantitation assay

Actin polymerization assay was done on WT (n=6) and BSMKO (n=7) aortas, according to Kim et al.^19^ with slight modifications. Briefly, each aortic strip (∼9 mg wet weight) was placed in homogenization tubes containing 120 μl of pre-warmed lysis and F-actin stabilization buffer (50 mM PIPES pH 6.9, 50 mM NaCl, 5 mM MgCl_2_, 5 mM EGTA, 5%(vol/vol) glycerol, 0.1% Nonidet P40, 0.1% Triton X-100, 0.1% Tween 20, 0.1% 2-mercapto-ethanol, 0.001% AntifoamC with 1 mM ATP and protease inhibitors) and homogenized in a Precellys 24 tissue homogenizer (Bertin Technologies, Rockville, MD, USA). Tissue homogenates were collected and incubated at 37°C for 20 minutes prior to ultracentrifugation. The homogenates were transferred to an optima TLX ultracentrifuge (Beckman Coulter, Brea, CA, USA) and spun at 150,000 g for 1 hour at 37°C. The supernatants (G-actin fractions) were collected and stored on ice. The pellets (F-actin fraction) were resuspended in 120 μl F-actin depolymerization buffer (8M Urea) on ice, by trituration every 15 minutes, for 1 hour to dissociate F-actin. Resuspended0020solutions were spun at 2,300 g for 5 min at 4°C to collect F-actin fraction (supernatant). Each sample was diluted with appropriate loading buffer and heated for 2 minutes at 95°C. Equal volume of samples were loaded on SDS-PAGE for further analysis of G-actin and F-actin by Western Blotting with anti-actin antibody (catalog # AAN01, Cytoskeleton Inc., Denver, CO, USA) at 1:500 dilution and IRDye 680RD anti-rabbit secondary antibody (catalog # 926-68071, LiCOR, Lincoln, NE, USA) at 1:10,000 dilution. The ratio of F-to G-actin was determined by scanning band densitometry on the raw images obtained using the Odyssey infrared imaging system (LiCOR).

### Statistical analysis

All data are expressed as mean ± SEM and were analyzed with GraphPrism software (v.7). *Ex vivo* aortic ring isometric tension measurements were analyzed by unpaired parametric test with Welch’s corrections. Repeated measures two-way ANOVA with Bonferroni’s multiple comparisons post-hoc test was used to analyze blood pressure over time in WT and BSMKO mice, at baseline and after angiotensin II administration. PWV in WT and BSMKO mice were analyzed by unpaired Student’s t-test. For Western blots and qRT-PCR results non-parametric Whitney-Mann t test was used. P values < 0.05 were considered significant.

### Data availability

Data supporting the findings of this study are available within the article and its Supplementary Information files and from the corresponding author upon request.

## RESULTS

### Genetic locus with genome-wide association with arterial stiffness

DB129663 is a 542bp non-coding transcript (PhyloSCF non-coding score based on small open reading frame analysis is - 173.3629; transcripts with a score < 41 are considered non-coding ^20^, LNCipedia) that partially overlaps a *BCL11B* enhancer in the genetic locus on chromosome 14 with genome-wide association with elevated PWV ^1^. A detailed analysis revealed that DB129663 gene contains a highly conserved 550bp sequence (> 95% homology among species), where the highest significant SNP variant associated with increased PWV (rs1381289) is located. By using BLAST, we located this highly conserved 550bp sequence in the mouse genome and identified AI060616 as a putative DB129663 homolog in the mouse genome. Considering that enhancer-associated lncRNAs are generally transcribed from the active enhancer they overlap ^21^, we used DB129663/AI060616 expression as a surrogate marker of a functionally active *BCL11B* enhancer in aortic tissue. We detected, for the first time, AI060616 in murine aortas as indicated by one specific PCR band of expected 356 bp molecular size, confirmed as AI060616 by sequencing, in aortic RNA extracts (Figure 2A). Similarly, DB129663 was detected in human aortas by qRT-PCR (Figure 2B). Notably, DB129663 knockdown with siRNA in cultured cells (Figure 2C) resulted in ∼ 50% decrease in Bcl11b mRNA. Overall, our results are consistent with the hypothesis that DB129663 is co-regulated with the 3’-*BCL11B* enhancer to transcriptionally regulate *BCL11B* expression.

**Figure 2.**
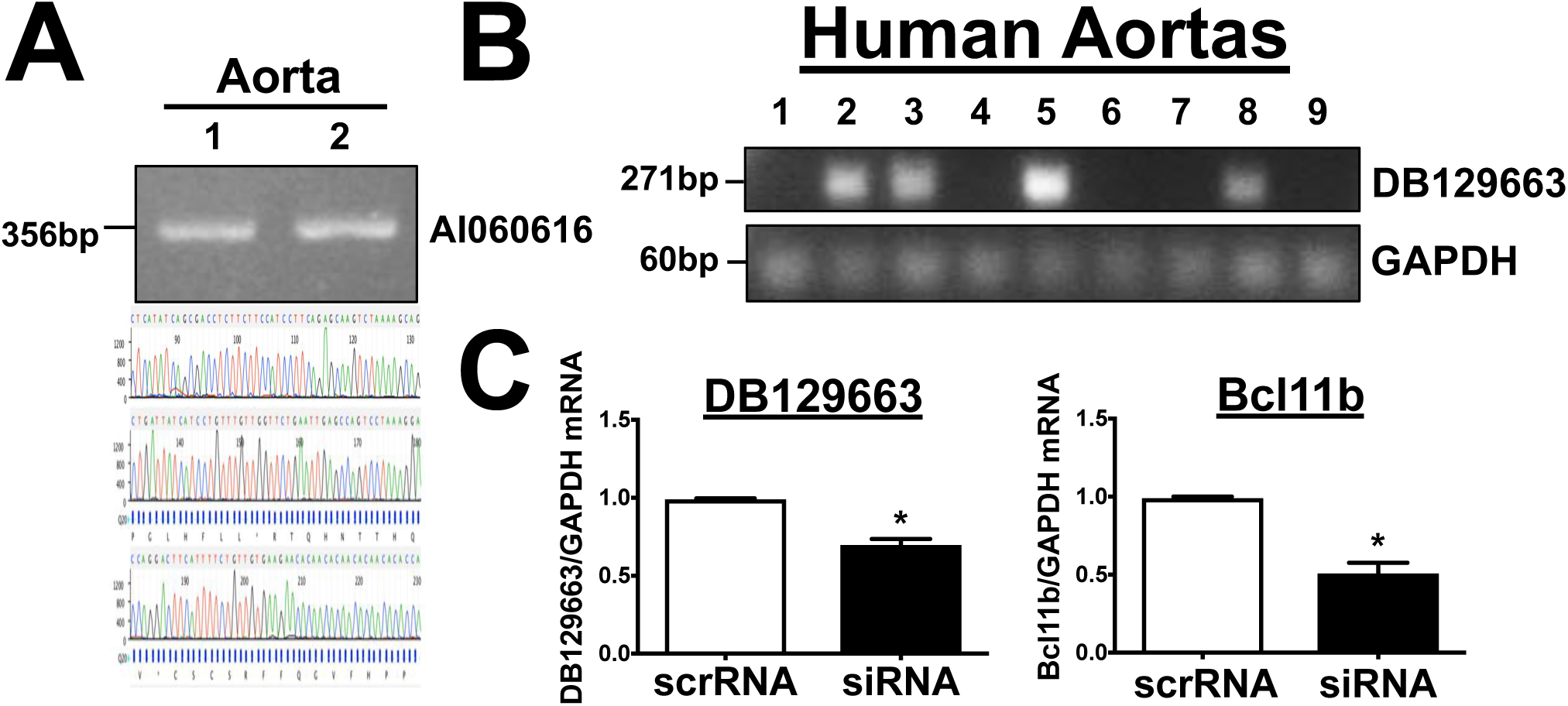
LncRNA DB129663 is expressed in the aorta. **(A)** LncRNA AI060616, the putative mouse homolog of DB129663, is expressed in murine aortas. AI060616 356bp nucleotide sequence was confirmed by sequencing PCR products from aortic RNA/cDNA, obtained from two C57Bl/6J mice (1 and 2). Sequencing chromatogram tracing shown in lower panel. **(B)** DB129663 is variably expressed in human aortas, as shown in 1% agarose gel electrophoresis of qRT-PCR products of aortic RNA/cDNA from 9 human subjects. Expected band sizes: 271 and 60bp for DB129663 and GAPDH, respectively. GAPDH serves as endogenous housekeeping gene for qRT-PCR. Each lane represents one human subject. **(C)** DB129663 silencing with siRNA (*left*) decreased Bcl11b expression in HEK cells (*right*), compared to non-targeting (scrambled) RNA (scrRNA), as measured by qRT-PCR. n=4. *, p < 0.05.

### Bcl11b is expressed in the VSM and is down-regulated in animal models of arterial stiffness

We next sought to determine whether Bcl11b, a downstream target of the DB129663/enhancer locus with SNP variants associated with arterial stiffness, is present in the vasculature. By using knock-in mice expressing a red fluorescence protein (mTomato) upon removal of *BCL11B* after tamoxifen administration (ER-Cre-Bcl11b^flox/flox^-mT knock in mice), we were able to visualize Bcl11b’s localization in aortic sections and, specifically, in the tunica media (Figure 3A, left panels). These findings were confirmed by immunostaining aortic sections with an antibody specific to Bcl11b (Figure 3A, right panels), with Western blotting of aortas and aortic smooth muscle cells (Figure 3B) and with qRT-PCR of human aortas (Figure 3C). In addition to the aorta, Bcl11b was visualized in VSM of arteries and arterioles in heart, lung and kidney (Supplemental Figure 1). Moreover, Bcl11b mRNA and protein (Figures 4A & 4B) levels were significantly decreased in aortas of high-fat, high-sucrose (HFHS)-fed obese mice, our established model of arterial stiffness ^11,13^, in comparison to normal diet (ND)-fed mice (WB band intensity: 1.00 ± 0.17 A.U. in ND, n=6 vs 0.33 ± 0.10 A.U. in HFHS, n=6; *, p<0.05). Similarly, Bcl11b mRNA levels were significantly decreased in aortas of 29-month-old mice, another established model of arterial stiffness, compared to 5-month-old-mice (Figure 4A; Bcl11b mRNA normalized to β-actin mRNA: 0.99 ± 0.01 in young, n=4 vs 0.08 ± 0.06 in old, n=4; *, p<0.05).

**Figure 3.**
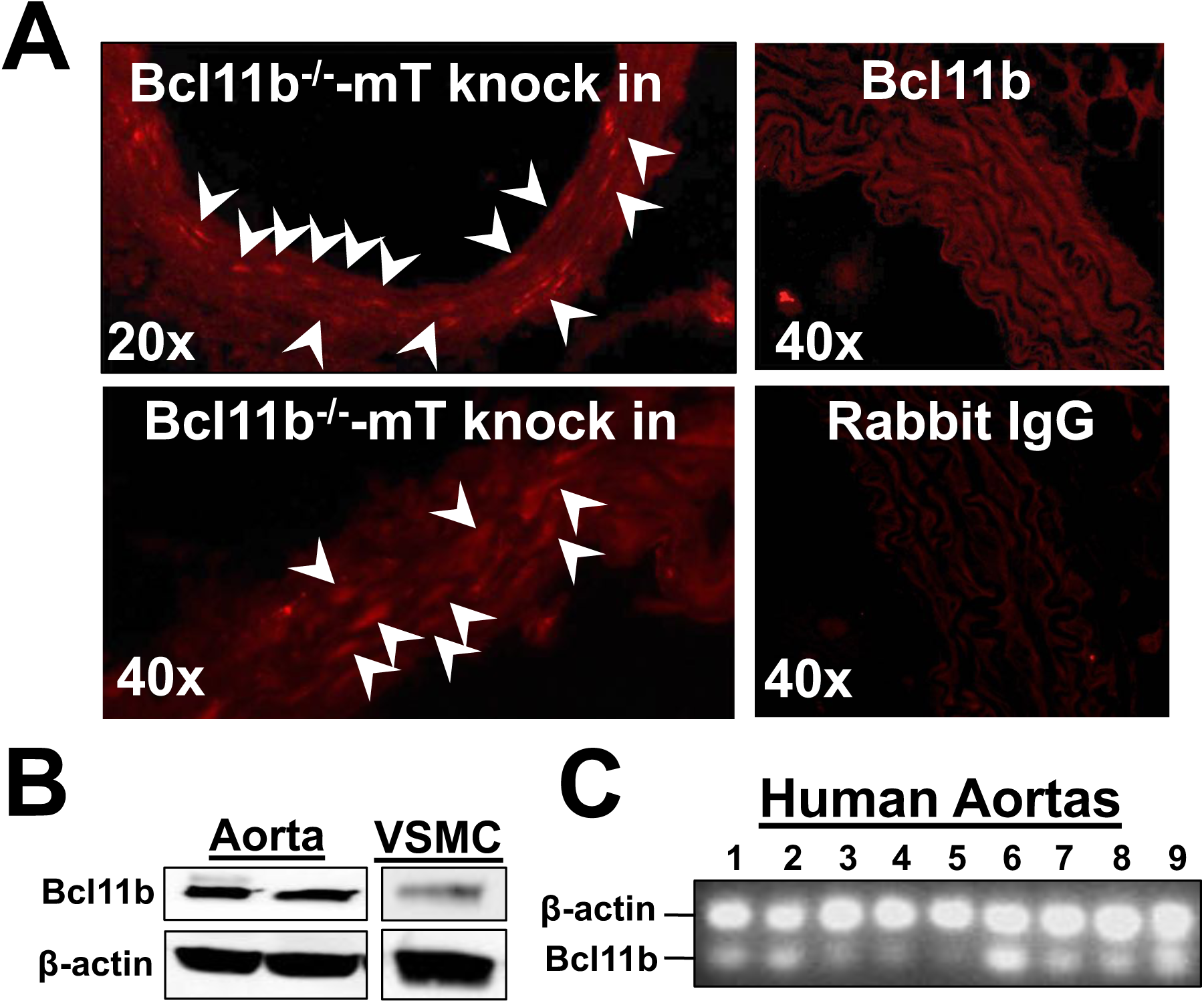
Bcl11b is expressed in vascular smooth muscle (VSM). **(A)** Representative fluorescent images of aortic sections (*top left panel*: 20X magnification; *bottom left panel:* 40X magnification) indicating Bcl11b’s localization in VSM. Red fluorescence indicates Tomato (mT) expression in lieu of Bcl11b, upon tamoxifen-induced Bcl11b removal, in ER-Cre-Bcl11bflox/flox-mT knock in mice. Arrowheads indicate clusters of VSM cells with high mT/Bcl11b fluorescence intensity. *Top right panel*: Representative image (40X magnification) of aortic section’s immunostaining with anti-Bcl11b IgG confirming Bcl11b localization in VSM. *Bottom right panel*: Incubation with rabbit IgG instead of primary antibody serves as negative control for antibody specificity. **(B)** Western blot on murine aortas and cultured aortic smooth muscle cells confirmed Bcl11b protein expression, with a band of expected MW 120kDa. β-actin serves as loading control. Each lane represents one mouse. **(C)** Image of 1% agarose gel electrophoresis of qRT-PCR Taqman products indicates Bcl11b mRNA expression in human aortas (n=9). Each lane represents one human subject. β-actin used as endogenous housekeeping gene in multiplexed assay.

**Figure 4.**
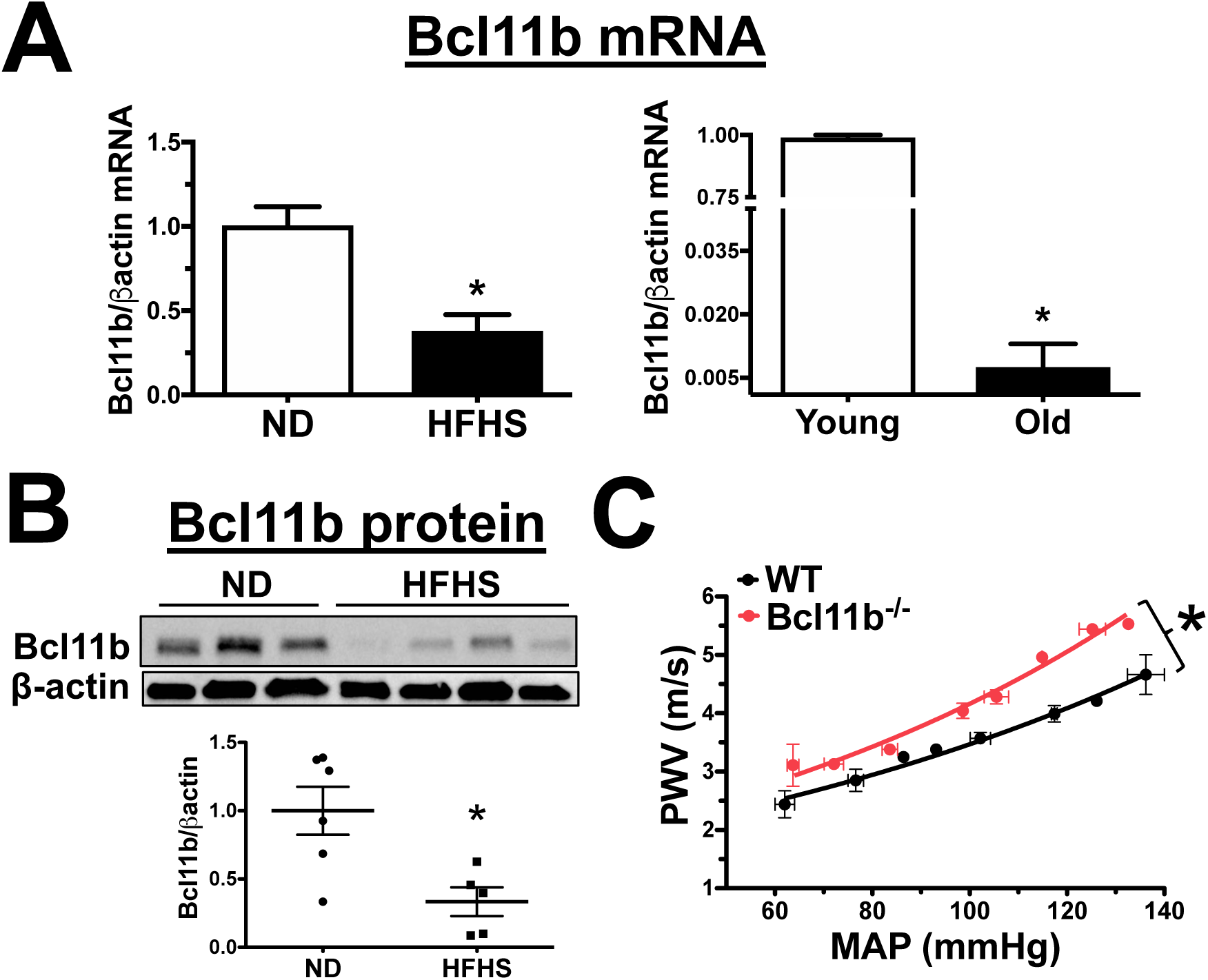
Bcl11b downregulation is associated with increased arterial stiffness. **(A)** Bcl11b mRNA levels measured by qRT-PCR are decreased in aortas of HFHS-fed obese mice (ND, normal diet; HFHS, high fat, high sucrose diet) and in aortas of 29-old mice (Old), two models of arterial stiffness, compared to ND-fed or 5 months-old (Young) mice, respectively. n=4 mice in each group; *, p < 0.05 ND vs HFHS or Young vs Old. **(B)** Representative Western Blot of Bcl11b protein expression in aortic homogenates of ND- and HFHS-fed mice. Each lane represents one mouse. β-actin serves as loading control. Scatter gram summarizes protein band quantitation (ratio of Bcl11b over β-actin band intensities), each dot represents one mouse. n=6 mice in each group; *, p < 0.05. **(C)** Pulse wave velocity (PWV, m/s), the in vivo index of arterial stiffness, measured over a range of mean arterial pressures (MAP, mmHg), is increased in 10-month old mice lacking Bcl11b compared to WT littermates. n=4 mice in each group; area under the curve: 260.3 m/s*mmHg in WT vs 285.6 m/s*mmHg in Bcl11b-/-; *, p < 0.05.

To further elucidate a functional role of vascular Bcl11b *in vivo*, we measured PWV in mice with global Bcl11b knock out (ER-Cre-Bcl11b^flox/flox^ treated with tamoxifen). We found that PWV was significantly increased in 10-month-old Bcl11b null mice compared with WT littermate controls (area under the curve: 260.3 m/s*mmHg in WT vs 285.6 m/s*mmHg in Bcl11b^-/-^, p < 0.05; over a range of mean arterial pressures, as shown in Figure 4C). Taken together, our novel findings demonstrate that Bcl11b is present in the VSM of the aortic wall and that aortic Bcl11b down-regulation may increase aortic stiffness.

### Bcl11b expression directly correlates with VSM contractile protein expression

Previous studies by us ^11,14,18^ and others ^22,23^ demonstrated that the maintenance of a contractile VSM cell phenotype is crucial in sustaining aortic wall mechanics and preserving arterial compliance. Based on the novel findings that Bcl11b is expressed in VSM and its decreased aortic levels are associated with increased arterial stiffness, we sought to determine whether vascular Bcl11b regulates arterial stiffness by regulating VSM contractile phenotype and function. We first compared aortic and VSM cell levels of Bcl11b and markers of VSM contractile phenotype, namely contractile proteins smooth muscle myosin heavy chain (gene ID *MYH11*), smooth muscle α-actin (α-SMA, gene ID *ACTA2*) and myocardin (gene ID *MYOCD*), a nuclear transcription factor and major transcriptional regulator of *MYH11* and *ACTA2* expression. VSM contractile phenotype is lost when VSM cells are isolated from the aorta and placed in culture, as they acquire a de-differentiated and proliferative phenotype. As expected, MYH11, α-SMA and MYOCD mRNA were significantly decreased in cultured VSM cells compared with aortas as was Bcl11b mRNA (Figure 5A). Removing fetal bovine serum (FBS) from culture medium, known to partially restore cultured VSM cells to a more differentiated state ^24^, increased Bcl11b and VSM contractile markers expression in VSM cells (Figure 5B). Our data suggest that dynamic changes in Bcl11b levels in VSM cells may be important in maintaining VSM cells in a differentiated and contractile state.

**Figure 5.**
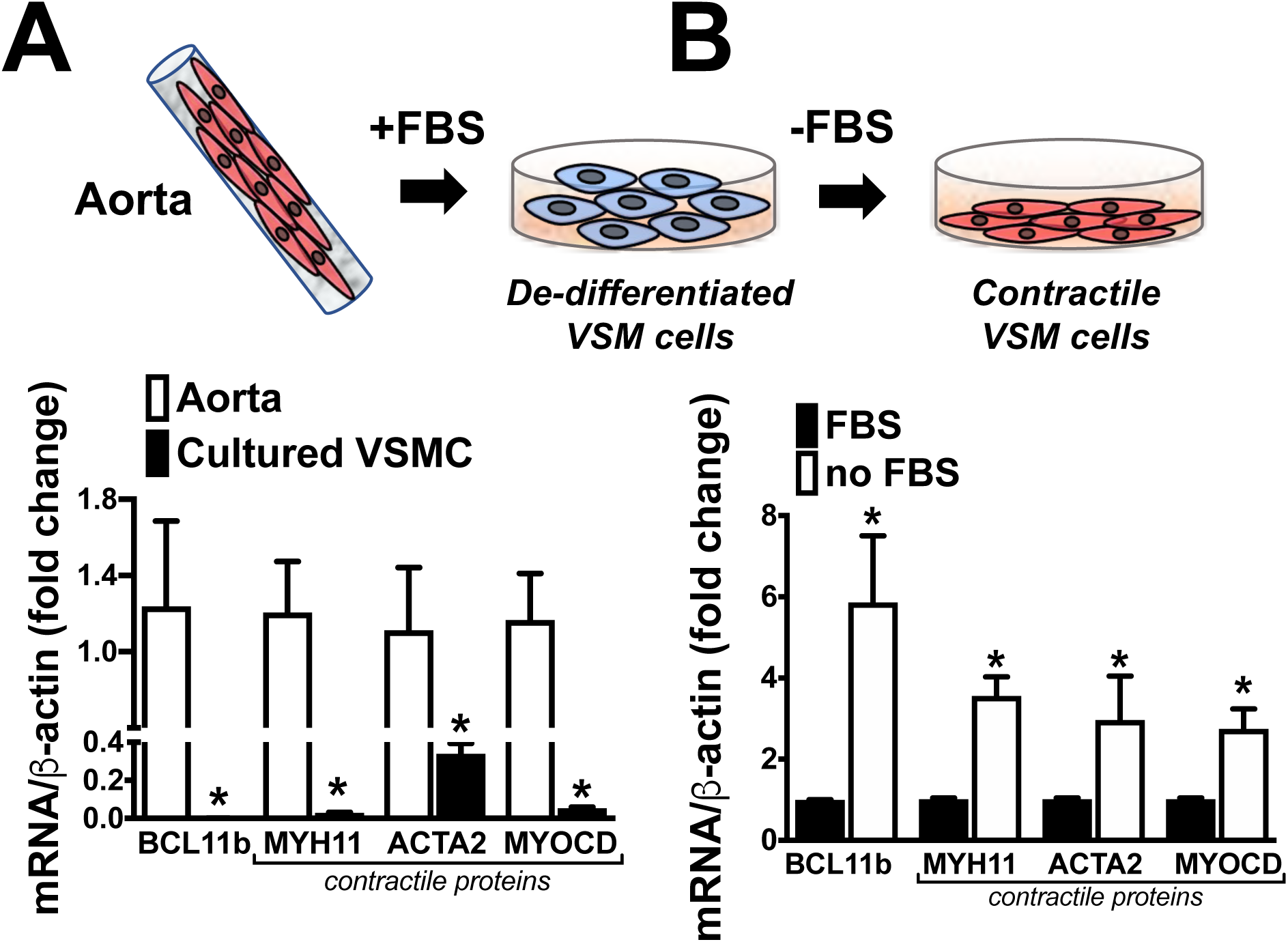
Bcl11b expression correlates with VSM contractile markers. **(A)** mRNA levels of Bcl11b and three markers of differentiated/contractile VSM cells, smooth muscle myosin *(MYH11),* smooth muscle a-actin *(ACTA2)* and myocardin *(MYOCD),* measured by qRT-PCR, are decreased in de-differentiated VSM cells cultured in presence of FBS, compared to aorta. 500ng of cDNA, prepared from aortic or VSM cell mRNA was used as input for qRT-PCR. B-actin used as endogenous housekeeping gene. n=5 replicate experiments. *, p < 0.05 aorta vs cultured VSM cells. **(B)** mRNA levels of Bcl11b, MYH11, ACTA2 and MYOCD are increased when cultured VSM cells are partially reversed to a more contractile phenotype by culturing them for 72 hrs in absence of fetal bovine serum (FBS). n=4 replicate experiments. *, p < 0.05, FBS vs no FBS.

### VSM Bcl11b transcriptionally regulates VSM contractile protein expression

To determine a direct role of Bcl11b in VSM phenotypic modulation, we generated mice with tamoxifen-inducible Bcl11b deletion in VSM (BSMKO; Figure 6A). As before, FBS removal from culture medium partially restored MYH11 in WT VSM cells but not in VSM cells isolated from aortas of BSMKO mice. Whether in presence or absence of FBS, lack of VSM Bcl11b was associated with a remarkable decrease in MYH11 mRNA (Figure 6B) and protein levels (Figure 6C; WB band intensity: 1.00 ± 0.05 A.U. in WT vs 0.42 ± 0.04 A.U. in BSMKO, n=5 replicate experiments; *, p<0.05) and α-SMA (Figure 6C; WB band intensity: 1.00 ± 0.04 A.U. in WT vs 0.63 ± 0.06 A. U. in BSMKO, n=4 replicate experiments; *, p<0.05). FBS removal or overnight incubation of FBS-starved VSM cells with TGFβ (20 ng/ml), a stimulus for α-SMA mRNA expression, increased α-SMA in WT VSM cells but to a significantly lower extent in BSMKO cells (Figures 6D & Supplemental Figure 2A). Removal of FBS decreased ki67 mRNA, a marker of cell proliferation, in both WT and BSMKO cells, as expected in quiescent cells; the decrease was more profound in WT than in BSMKO VSM cells (Supplemental Figure 2B). Our findings of decreased MYH11 and α-SMA were corroborated in a second animal model in which constitutive Bcl11b removal in VSM was achieved with a SM22α (transgelin) promoter-driven Cre recombinase transgene (Supplemental Figure 2C). Overall, our data indicate that Bcl11b is important to sustain the expression of VSM-specific contractile proteins MYH11 and α-SMA in order to preserve VSM cells in a differentiated and quiescent state.

**Figure 6.**
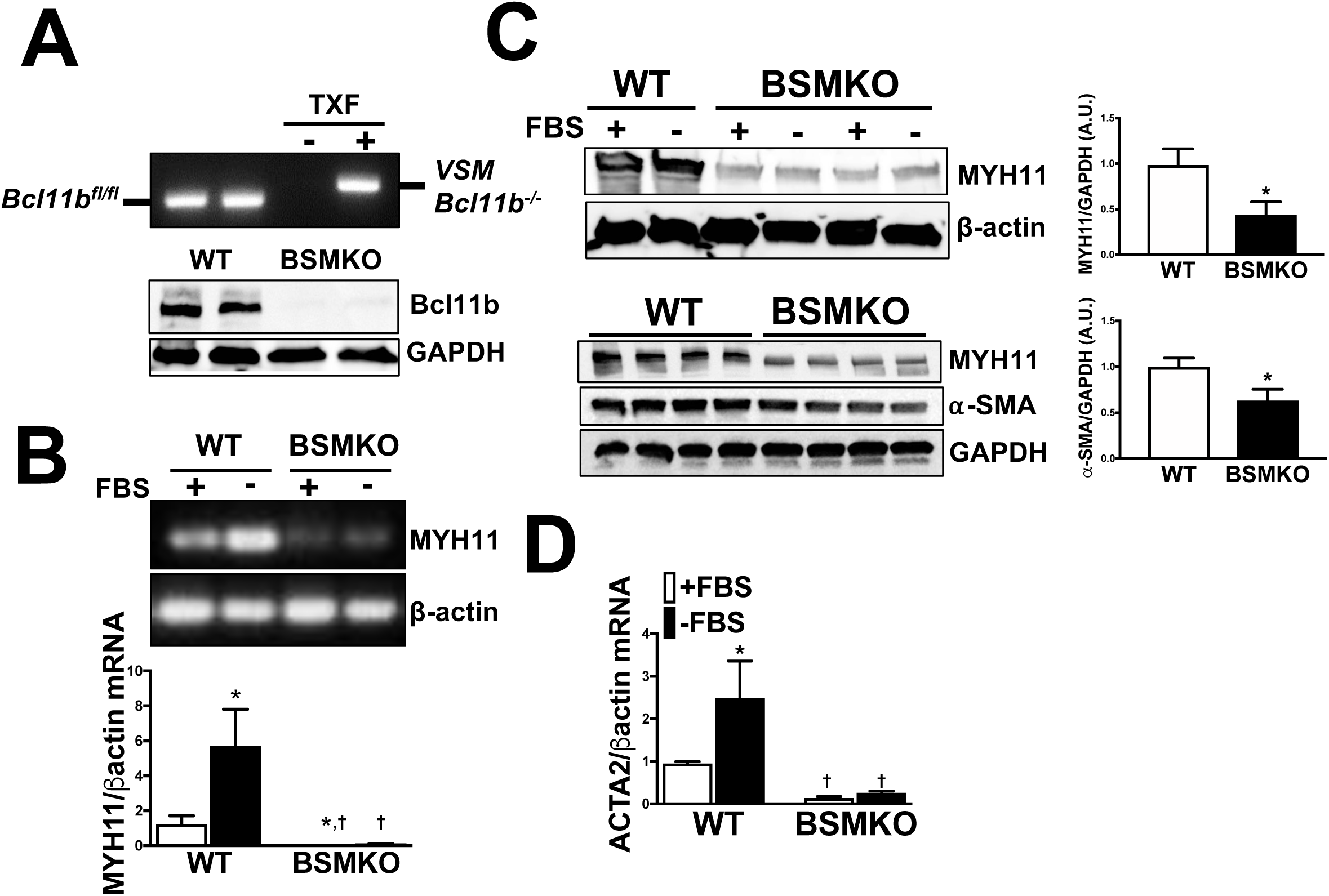
VSM Bcl11b transcriptionally regulates VSM contractile proteins. **(A)** *Top.* Representative image of genomic DNA PCR products from MYH11-ER-Cre+ floxed Bcl11b mice, before and after vehicle or tamoxifen (TXF) administration (expected band size: 350bp for Bcl11bfl/fl; 450bp for VSM Bcl11b-/-). *Bottom.* Western blot of aortic homogenates from 2 MYH11-ER-Cre+ floxed Bcl11b mice receiving vehicle (WT) and 2 MYH11-ER-Cre+ floxed Bcl11b mice receiving tamoxifen (BSMKO), confirming VSM-specific Bcl11b deletion. The expected band of molecular weight 120kDa is visible in WT aortic media (adventitia removed) but not in VSM-specific BSMKO. GAPDH serves as loading control. **(B)** qRT-PCR products for MYH11 and housekeeping gene β-actin shown on 1% agarose electrophoresis gel. Graph summarizes MYH11 mRNA qRT-PCR quantitation, calculated with the δδCt method, in VSM cells isolated from aortas of BSMKO mice relative to WT littermates. VSM cells were cultured in presence or absence of fetal bovine serum (FBS), a regulator of contractile proteins in VSM cells. n=5 replicate experiments. *, p < 0.05 FBS vs no FBS;, p < 0.05 WT vs BSMKO. **(C)** Representative Western Blot demonstrating decreased smooth muscle myosin (MYH11) and smooth muscle α-actin (α-SMA) expression in BSMKO VSM cells, cultured with or without FBS compared to WT. Band intensity quantitation summarized in graph. n=5 replicate experiments, *, p < 0.05 WT vs BSMKO. **(D)** Smooth muscle α -actin (*ACTA2)* mRNA levels were decreased in BSMKO VSM cells cultured in presence or absence of FBS, compared to WT cells. n=4 replicate experiments. *, p < 0.05 FBS vs no FBS;, p < 0.05 WT vs BSMKO.

### Bcl11b regulates VSM tone and stiffness

In order to examined whether VSM-specific Bcl11b-dependent decreases in contractile filament components (MYH11 and α-SMA) were associated with functional consequences in the vasculature, we measured (1) force, wall tension and stress generated by WT and BSMKO aortic rings; (2) arterial stiffness *in vivo* by PWV in WT and BSMKO mice; and (3) blood pressure in WT and BSMKO mice, at baseline and after angiotensin II administration. Bcl11b deletion in VSM did not affect gross aortic morphology, as indicated by comparable aortic media thickness (54.5 ± 1.0 μm in WT, n=5 vs 55.1 ± 1.1 μm in BSMKO, n = NS) and diameter (unloaded dimensions: 0.74 ± 0.02 mm in WT, n=5 vs 0.75 ± 0.01 mm in BSMKO, n=7; NS) between WT and BSMKO mice (Supplemental Figure 3). However, baseline force (1010 ± 96 mg in WT, n=5 vs 1511 ± 106 mg in BSMKO, n=7; *, p<0.05), wall tension (1.72 ± 0.13 N/m in WT, n=5 vs 2.60 ± 0.24 N/m in BSMKO, n=7; *, p<0.05) and stress (17.7 ±1.5 kPa in WT, n=5 vs 25.9 ± 2.2 kPa in BSMKO, n=7; *, p<0.05) generated by BSMKO aortic rings in organ baths were significantly increased compared to WT (Figure 7A). Likewise, aortic stiffness measured *in vivo* by PWV was significantly increased in BSMKO mice 2 months after VSM Bcl11b removal, compared to littermate WT controls (Figure 7B; 3.1 ± 0.1 m/s in WT, n=14 vs 3.8 ± 0.2 m/s in BSMKO, n=17; *, p<0.05). No differences in PWV increases were observed between BSMKO (4.0 ± 0.4 m/s, n=9; 2 months after tamoxifen administration) and SM22BKO mice (3.5 ± 0.1 m/s, n=8; 2 months of age; p=0.3) nor between male (3.9 ± 0.4 m/s, n=12) and female mice (3.5 ± 0.2 m/s, n=5; p=0.6).

**Figure 7.**
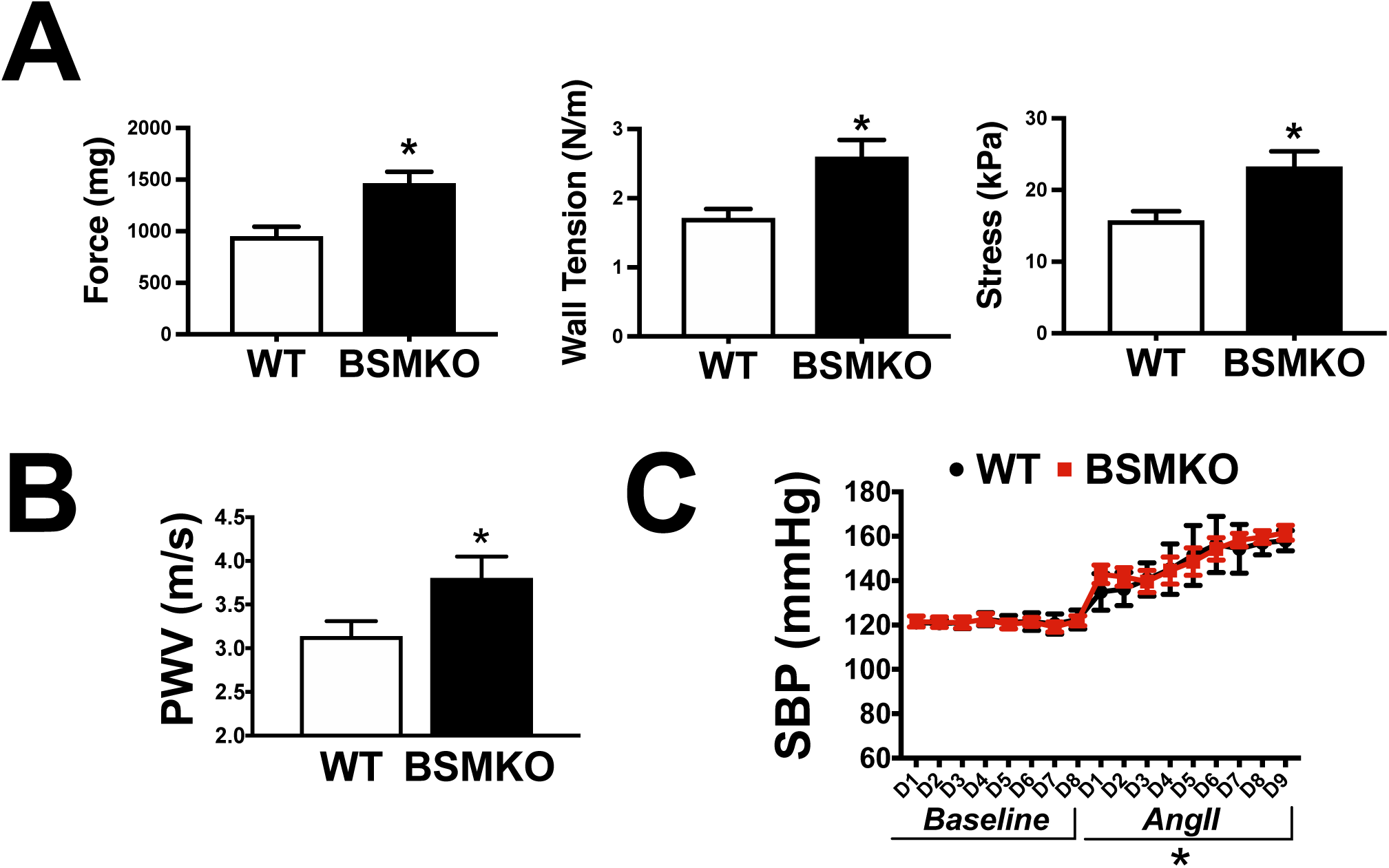
VSM Bcl11b regulates VSM tone and stiffness. **(A)** Force (mg), wall tension (N/m) and stress (kPa) measured *ex vivo* in WT (n=5) and BSMKO (n=7) aortic rings. *, p < 0.05. **(B)** BSMKO mice (n=17) have increased arterial stiffness, measured in vivo by PWV (m/s), compared to WT littermate controls (n=14). *, p<0.05. **(C)** Radiotelemetry blood pressure measurements in WT (n=7) and BSMKO (n=8) mice before and after angII administration (1 mg/kg/d) by osmotic minipumps. SBP, systolic blood pressure. *, p<0.05, D1 to D9 angII vs D1 to D8 baseline by repeated measures two-way ANOVA for both WT and BSMKO.

Bcl11b deletion in VSM did not significantly affect baseline blood pressure (systolic blood pressure, SBP: 121.8 ± 3.8 mmHg in WT, n=7 vs 120.0 ± 2.3 mmHg in BSMKO, n=8; NS) nor the development of angiotensin II-hypertension in 2-month old BSMKO mice (SBP: 156.5 ± 7.6 mmHg in WT, n=7 vs 151.7 ± 2.7 mmHg in BSMKO, n=8; NS) compared to age-matched WT littermate controls (Figure 7C and mean arterial pressure, MAP and diastolic blood pressure, DBP in Supplemental Figure 4A). However, two-week ang II treatment was sufficient to cause aortic aneurysms in BSMKO mice (5 out of 7, ∼ 70% incidence and 2 out of 7 died due to aneurysm rupture) in suprarenal and/or thoracic aortic regions, compared to angII-treated WT mice, none of which developed aortic dilatations in the same period (0 out of 7; Figure 8A). TUNEL staining of aortic sections indicated an increased number of apoptotic VSM cells in aneurysmal aortas of angII-treated BSMKO compared to angII-treated WT mice (Figure 8B).

**Figure 8.**
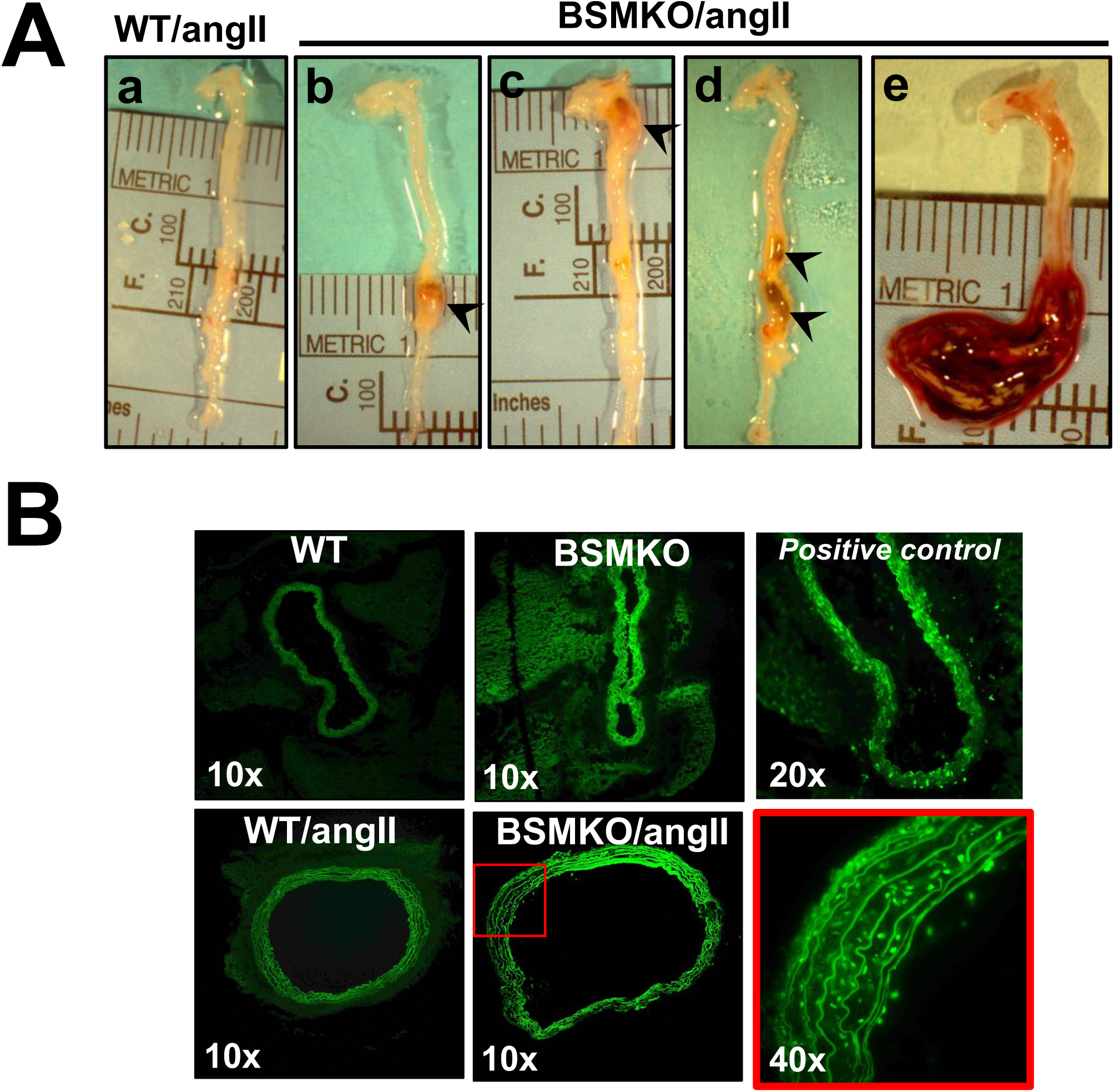
VSM Bcl11b null mice develop aortic aneurysms in response to angiotensin II. **(A)** Representative images of WT and BSMKO aortas after 2-week treatment with angII. 70% of BSMKO mice developed aortic aneurysm in the suprarenal **(b)** or thoracic **(c)** region or both **(d)** (indicated by arrowheads), while no aneurysm developed in angII-treated WT mice **(a)**. **(e)** aorta from an angII-treated BSMKO mouse found dead. Metric ruler shown as reference for microscope magnification. **(B)** Representative images of TUNEL staining of aortic sections from control and angII-treated WT and BSMKO mice indicating increased number of apoptotic VSM cells in angII-treated BSMKO aortas (shown at increased magnification in *red box*). FITC-labeled nuclei shown as fluorescent green dots indicate apoptotic cells. Positive control is a WT aorta pre-treated for 30 min with Dnase I, as described in Methods. Magnification indicated on image.

### VSM Bcl11b regulates actin polymerization and vasodilator-stimulated phosphoprotein (VASP) phosphorylation at serine 239

Non-muscle actin polymerization in the cytoskeleton contributes to VSM cell cytoskeletal stiffness ^25^ and VSM tension development independently of myosin light chain 20 (MCL_20_) phosphorylation and cross-bridge cycles ^25^. Therefore, we next examined whether cytoskeletal actin polymerization may have contributed to increased VSM tone and stiffness in BSMKO aortas despite decreases in contractile proteins. We found increased filamentous to glomerular (F/G) actin ratio, indicative of increased actin polymerization, in BSMKO aortas compared to WT, assessed by Western Blot in aortic homogenates (Figure 9A; ratio of F and G actin bands intensity: 4.82 ± 1.33 A.U. in WT, n=6 vs 9.09 ± 1.02 A.U. in BSMKO, n=7; *, p<0.05).

**Figure 9.**
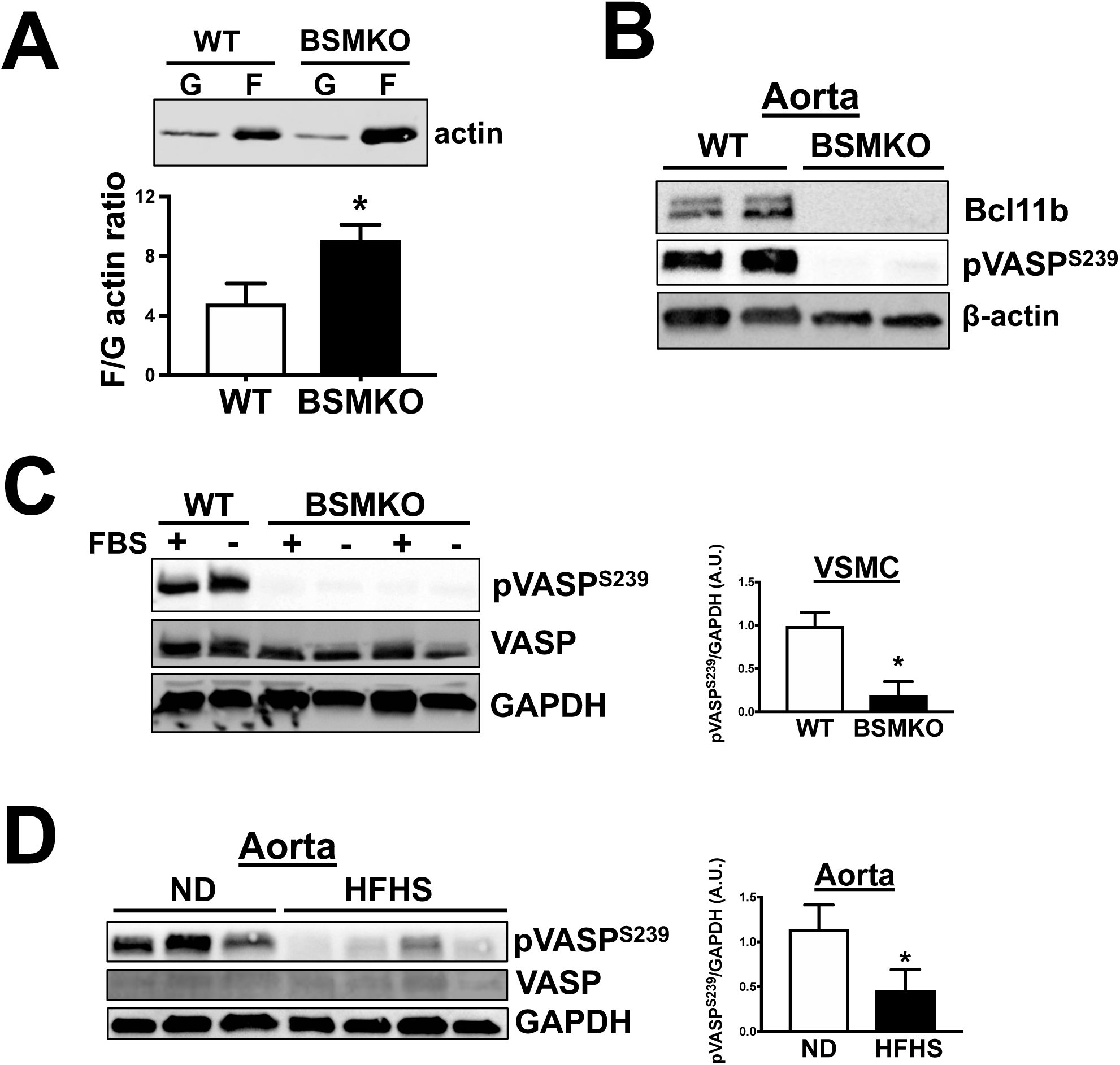
VSM Bcl11b regulates pVASPS239. **(A)** Representative Western Blot images of F and G actin in WT (n=6) and BSMKO (n=7) aortas. F/G actin ratio quantitation in graph. *, p < 0.05. **(B)** Representative Western Blot images demonstrating impaired VASP phosphorylation at serine 239 (pVASPS239) in aortas of BSMKO mice (n=4) compared to WT littermate controls (n=4). β-actin serves as loading control. Each lane represents one mouse. **(C)** pVASPS239 was significantly decreased in BSMKO VSM cells, cultured with or without FBS, compared to WT cells. Total VASP remained unchanged. GAPDH serves as loading control. n=5 replicate experiments. **(D)** pVASPS239 was significantly decreased in aortas of HFHS-fed mice (n=10) compared to ND-fed controls (n=6). Total VASP was similar in the two groups. GAPDH serves as loading control. Each lane represents one mouse. Band intensity quantitation summarized in graphs. *, p < 0.05 WT vs BSMKO or ND vs HFHS.

Vasodilator-stimulated phosphoprotein (VASP) is an important regulator of non-muscle actin polymerization-dependent VSM tone ^26^ while phosphorylation of VASP at serine 239 (pVASP^S239^) inhibits actin polymerization in VSM cells ^27^. Based on our previous findings that increased aortic pVASP^S239^ is associated with decreased diet-induced arterial stiffness ^11^, we next examined whether VASP and pVASP^S239^ were affected in VSM Bcl11b-deleted aortas thereby regulating cytoskeletal actin assembly. We found that pVASP^S239^ was dramatically decreased in BSMKO compared to WT aortas (Figure 9B), and in BSMKO VSM cells compared to WT cells, cultured with or without FBS (Figure 9C; WB band intensity: 0.99 ± 0.04 A.U. in WT vs 0.19 ±0.04 A.U. in BSMKO, n=5 replicate experiments; *, p<0.05), while total VASP remained unchanged. Similar findings were obtained in aortas of HFHS-fed obese mice (i.e., with decreased aortic Bcl11b, Figure 4B), compared to ND-fed mice (Figure 9D; WB band intensity: 1.14 ± 0.11 A.U. in ND, n=6 vs 0.46 ± 0.07 A.U. in HFHS, n=10; *, p<0.05).

VASP phosphorylation in VSM cells is finely regulated by protein kinases (PKG, PKA) ^28^ and phosphatases (PP1, PP2A, PP2B, PP2C) ^29^. Of interest, Ca^++^/calmodulin-dependent serine-threonine phosphatase calcineurin (PP2B) has been shown to directly interact with Bcl11b to regulate gene expression in T-cells ^30^. Therefore, we examined whether PP2B may be responsible for decreased pVASP ^S239^ levels in BSMKO VSM cells. We found that PP2B protein expression was significantly upregulated in BSMKO VSM cells (Figure 10A). In addition, overnight treatment with cyclosporine A, a PP2B inhibitor, restored pVASP ^S239^ levels in BSMKO VSM cells (Figures 10B & 10C; quantitation in graph), in a dose-dependent manner, indicating that decreased pVASP ^S239^ in BSMKO VSM is dependent mainly on increased PP2B. Moreover, overexpressing Bcl11b in aortic media rings by transient transfection for 3 days was sufficient to significantly restore MYH11, α-SMA, pVASP ^S239^ and decrease PP2B to control levels (Figure 10D). Taken together, our data indicate that VSM Bcl11b regulates PP2B-dependent decreases in pVASP^S239^ associated with increased actin polymerization.

**Figure 10.**
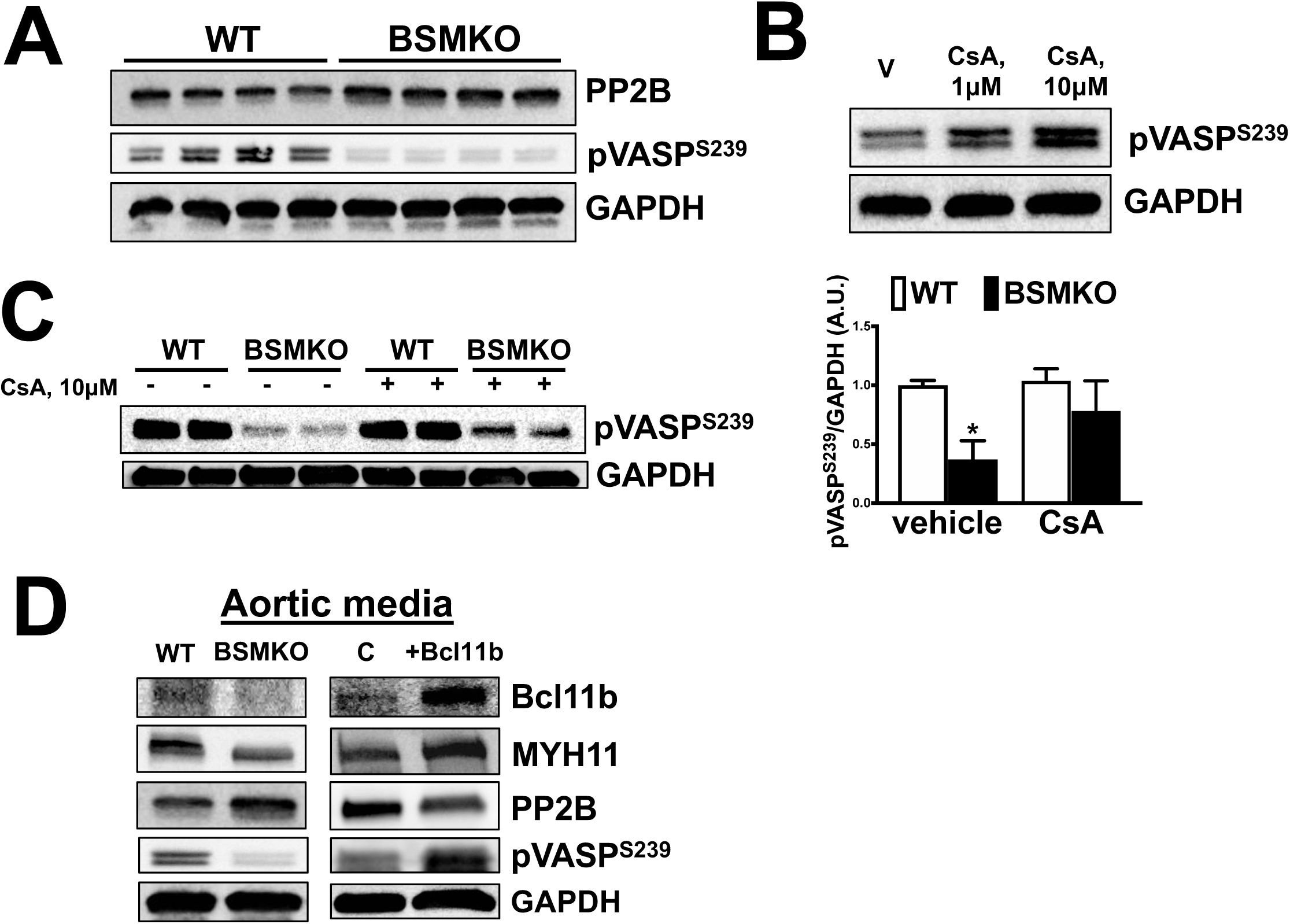
VSM Bcl11b regulates pVASPS239 via calcineurin. **(A)** Representative Western Blot images demonstrating increased calcineurin (PP2B) expression and impaired VASP phosphorylation at serine 239 (pVASPS239) in BSMKO VSM cells compared to WT controls. GAPDH serves as loading control. Each lane represents a cell preparation from one mouse. Same protein homogenates and Western Blot as in Figure 6C. **(B&C)** Treatment of BSMKO VSM cells with 1 or 10 μM cyclosporine A (CsA), a calcineurin inhibitor, normalized pVASPS239 in a dose-dependent manner. V, vehicle control. Quantitation of band intensities in graph (n=3 replicate experiments). *, p < 0.05, WT/vehicle vs BSMKO/vehicle. **(D)** Western Blots for Bcl11b, MYH11, PP2B, pVASPS239 in WT and BSMKO aortic media (adventitia removed) without (*left*) or with (*right*) transient transfection with vehicle (Lipofectamine; C, control) or 20 μg Bcl11b plasmid. GAPDH serves as loading control.

## DISCUSSION

A recent genome-wide association study (GWAS) demonstrated that specific SNPs in the vicinity of the *BCL11B* genetic locus are associated with increased arterial stiffness and subsequent risk of developing CVD ^1^. Interestingly, the highest significant SNP variant (rs1381289) in this “aortic stiffness” locus falls in a highly conserved sequence within a lncRNA (DB129663) which partially overlaps with a *BCL11B* enhancer. These observations prompted us to examine whether there is a cause-effect relation between Bcl11b and vascular function.

Here we report for the first time that DB129663 and its putative mouse homolog AI060616 are expressed in human and murine aortas, respectively, and that DB129663 downregulation results in ∼ 50% decrease in Bcl11b mRNA, which is consistent with the hypothesis that the *BCL11B* enhancer region regulates Bcl11b expression at the transcriptional level. These findings suggest that SNP variants in the 3’-*BCL11B* locus may alter the Bcl11b gene enhancer function and may play a causal role in the pathogenesis of arterial stiffness by regulating Bcl11b expression.

Elevated PWV, the gold standard measure of aortic wall stiffness, strongly associates with adverse cardiovascular outcomes ^35–37^. Despite compelling clinical evidence, cellular and molecular cues of aortic wall stiffening are not fully understood, hampering the discovery of therapeutic targets that can slow or reverse arterial stiffness and thus decrease the risk of developing CVD.

Our study demonstrates for the first time that Bcl11b, previously known for its role in T lymphocyte ^8^ and neuronal ^9^ lineage commitment, is expressed in the VSM layer of the human and murine aortic wall and that VSM Bcl11b is a crucial regulator of VSM structural components and aortic stiffness, as corroborated by the following findings: (1) *in vivo* Bcl11b deletion in VSM (BSMKO mice) resulted in remarkable reductions in aortic smooth muscle myosin (MYH11) and smooth muscle α-actin (α-SMA) and increased non-muscle actin polymerization (F/G actin ratio); (2) mice lacking Bcl11b globally or specifically in VSM (BSMKO) have increased PWV, the *in vivo* index of arterial stiffness, compared to WT littermates; (3) aortic rings from BSMKO mice have increased baseline force, stress and wall tension compared to WT; and (4) Bcl11b is downregulated in aortas of high fat, high sucrose-fed obese mice and aged mice, two mouse models of arterial stiffness.

Interestingly, a major Bcl11b interacting protein, COUP-TFII, is required for atria and blood vessel development during embryogenesis ^38^. However, until now, a role of Bcl11b in the adult cardiovascular system was unknown. Our novel findings with a mTomato-tagged Bcl11b reporter mouse, further confirmed by qRT-PCR in human and murine aortas and Western Blotting and immunohistochemistry in murine aortas and VSM cells, clearly indicate that Bcl11b is expressed in the VSM of the aortic wall and in arteries and arterioles of heart, lung and kidney. Importantly, we uncovered a pivotal role of VSM Bcl11b in the regulation of VSM contractile and cytoskeletal filaments, which form a coordinated system to efficiently transduce contractile forces to the extracellular matrix and among adjacent VSM cells, thereby sustaining aortic wall mechanics and compliance.

In response to pressure or mechanical stretch, thin filament dynamic assembly, namely non-muscle actin polymerization, become a major determinant of VSM contraction and basal tone ^25,39^, independently of myosin light chain 2 phosphorylation and actino-myosin cross-bridges cycles ^40,41^. Dynamic actin cytoskeletal rearrangements can sustain VSM force generation and cytoskeletal stiffness to maintain vessel diameter in response to wall tension or stretch ^42^, as it may occur in the aorta exposed to cyclic strain induced by cardiac contraction, particularly in the proximal regions. Therefore, an increase in baseline filamentous/glomerular (F/G) actin ratio, indicative of increased actin polymerization (Figure 9A), is consistent with increased baseline force and wall tension in BSMKO aortas (Figure 7A) and increased PWV in BSMKO mice (Figure 7B) compared to WT controls.

Vasodilator-stimulated phosphoprotein (VASP) has emerged as an important mediator of actin polymerization-dependent VSM force generation. Specifically, we ^26^ and others ^43–45^ have previously shown that VASP interacts with α-actinin, vinculin, zyxin and other components of thin filament assembly at focal adhesion and dense bodies, which are important sites of VSM contractile filaments attachment, cell-cell interactions and cell adhesion to the extracellular matrix, thereby contributing to VSM tone and stiffness independently of myosin light chain 2 phosphorylation. The drug cytochalasin D, commonly used to block actin polymerization in a variety of cell types, is known to interfere with VASP localization to nascent F-actin filaments ^46^ underscoring the pivotal role of VASP in cytoskeletal actin rearrangement. Moreover, VASP overexpression has been shown to induce F-actin assembly ^47^; in contrast, VASP phosphorylation at serine 239 is sufficient to inhibit actin filament polymerization ^27^.

Based on these and our previous findings that increased aortic pVASP^S239^ is associated with decreased diet-induced arterial stiffness ^11^, we measured pVASP^S239^ and total VASP in aortas of mice lacking Bcl11b in VSM to assess whether these mechanisms could similarly occur in stiff aortas of BSMKO mice. We found that baseline pVASP^S239^ was dramatically decreased in BSMKO aortas and VSM cells, while total VASP remained unchanged. Moreover, in accordance with our previous work, stiff aortas of high fat, high sucrose-fed mice have decreased pVASP^S239^ levels which closely correlated with Bcl11b downregulation (Figures 4A, 4B & 9D).

Overall, our data suggest that VSM Bcl11b is required to regulate VASP-mediated actin polymerization and basal VSM tone in the aorta. Increased actin polymerization and associated increases in baseline wall tension and stiffness in BSMKO aortas, may be a compensatory mechanism required to sustain aortic wall mechanics in the settings of drastically decreased contractile elements MYH11 and α-SMA (see Figure 11 for a graphical summary).

**Figure 11.**
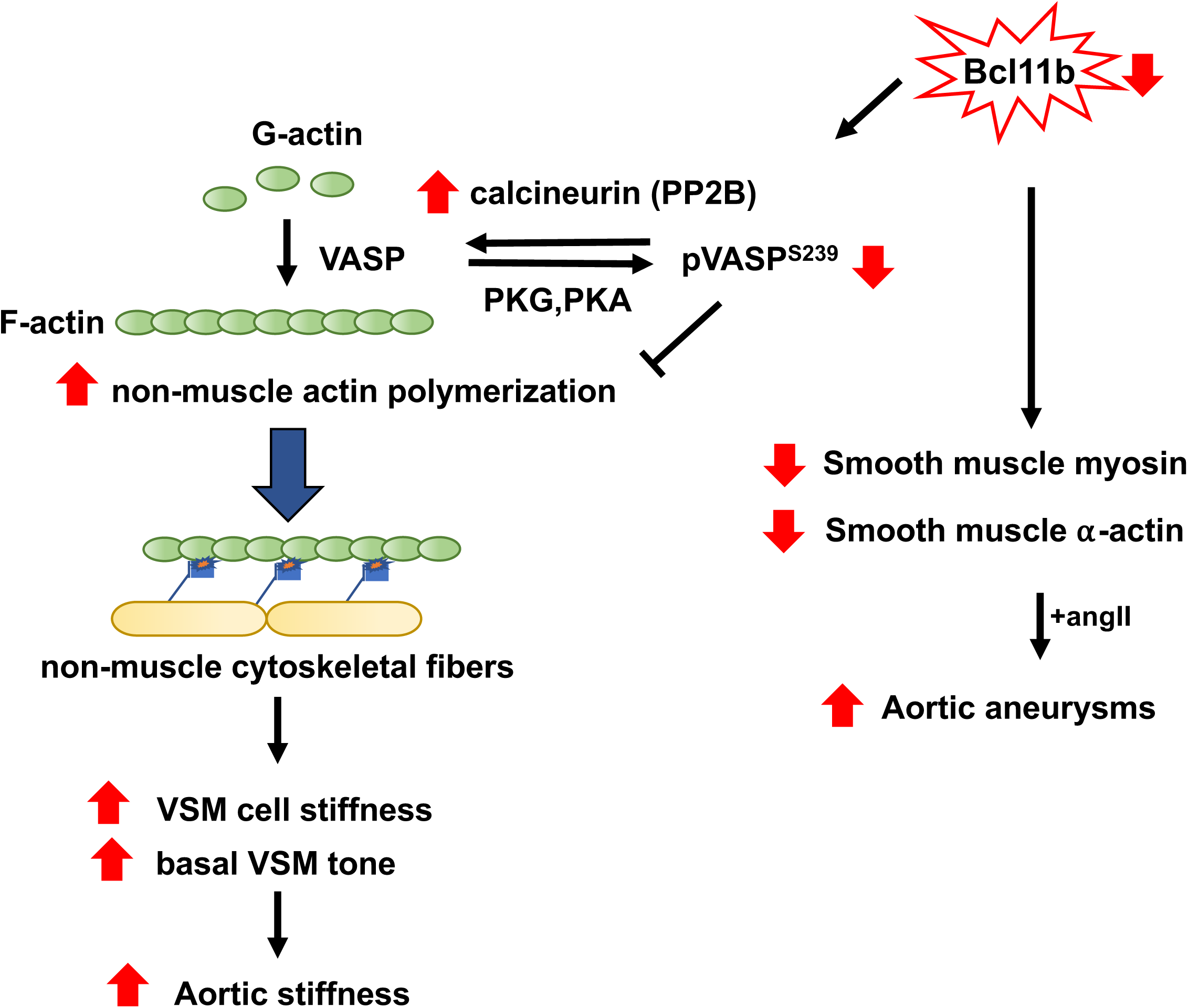
Postulated mechanisms by which VSM Bcl11b regulates vascular phenotype. Down-regulated Bcl11b in VSM decreases VASP phosphorylation at serine 239 (pVASPS239) via calcineurin (PP2B), resulting in increased F-actin polymerization, increased VSM cell stiffness and increased basal VSM tone, leading to increased aortic stiffness. In addition, down-regulated Bcl11b in VSM is associated with significant decreases in smooth muscle contractile proteins myosin and α-actin, which may contribute to increased incidence of aortic aneurysms in angiotensin II-treated BSMKO mice.

Considering that Bcl11b interacts directly with the Ca^++^/calmodulin serine-threonine phosphatase calcineurin (also known as protein phosphatase 2B, PP2B) in T-cells ^30^, we explored the possibility that a similar interaction could occur in VSM and may contribute to decreased pVASP^S239^ when VSM Bcl11b is deleted. Baseline PP2B levels were significantly increased and showed an inverse correlation with pVASP^S239^ in BSMKO VSM cells. Moreover, treatment with a calcineurin inhibitor, cyclosphorine A, was able to partially restore decreased pVASP^S239^ in BSMKO VSM cells. Although we did not determine the contribution of other possible kinases or phosphatases to the regulation of VASP phosphorylation in BSMKO aortas, our data strongly suggest that (1) VSM Bcl11b is a transcriptional repressor of PP2B in the aorta; and (2) Bcl11b-dependent PP2B is a primary regulator of VASP phosphorylation at serine 239 in VSM cells.

Bcl11b is a zinc finger protein known to bind G/C-rich DNA elements ^5^ some of which are negative-acting *cis*-elements crucial to regulate VSM de-differentiation ^48,49^. Therefore, it is reasonable to speculate that Bcl11b transcriptionally regulates MYH11, α-SMA and calcineurin by directly binding to their promoter regions, possibly by recruiting the histone deacetylase sirtuin-1 (SirT1) ^50^ to epigenetically regulate VSM gene promoters, although the detailed mechanisms remain to be determined. Our findings that transiently overexpressing Bcl11b in aorta for 3 days was sufficient to reverse the decrease in MYH11, α-SMA, pVASP^S239^ and the increase in calcineurin expression (Figure 10D) are consistent with this hypothesis.

Despite having profound effects on the aorta, lack of VSM Bcl11b did not affect blood pressure regulation in BSMKO mice as baseline and angiotensin II-hypertension were comparable with WT mice suggesting a dispensable role of Bcl11b in the regulation of VSM tone in resistance vessels. Mice were 2 months old at the time of BP measurements; therefore, we cannot exclude that detectable effects of VSM Bcl11b deletion on BP regulation may become apparent at later time points with advancing age. A differential role of Bcl1b in VSM of large *versus* small arteries could also be explained by the fact that smooth muscle along the adult vascular tree is not homogeneous but rather a mosaic of phenotypically and functionally distinct smooth muscle cell types ^54^. Elegant lineage mapping studies have demonstrated that smooth muscle of proximal regions of the aorta, which are the most susceptible to vascular stiffening, namely the outflow tract, innominate, common carotid and subclavian arteries, but not the smooth muscle of resistance vessels, developmentally originates from the cranial neural crest ^55^. Considering that Bcl11b is highly expressed and required for neuronal development during embryogenesis ^9,10^, it is reasonable to speculate that Bcl11b, possibly via COUP-TFII, may be a common regulator of lineage commitment for VSM and neurons in early embryonic development. Alternatively, a common progenitor neuronal stem cell expressing Bcl11b may be crucial for smooth muscle cell lineage commitment, including the expression of contractile markers, MYH11 and α-SMA, in large vessels during fetal development. *In vivo* lineage tracing studies, such as *LacZ* reporter mice, will be necessary to fully elucidate Bcl11b expression patterns during gestation and in the adult vascular tree.

Interestingly, despite the lack of effect of Bcl11b deletion in VSM on baseline blood pressure or ang II-hypertension, two week infusions with ang II were sufficient to induce thoracic or suprarenal aortic aneurysms in 5 out of 7 (∼ 70%) ang II-infused BSMKO mice, as compared to 28 days commonly required to observe suprarenal aortic aneurysms with ang II infusions in experimental models ^56^. MYH11 and α-SMA genetic mutations have been causally linked to non-syndromic aortic aneurysms ^57^. Considering that lack of VSM Bcl11b drastically reduced MYH11 and α-SMA expression in BSMKO mice (Figure 6), it is plausible that BSMKO mice phenocopy MYH11 and α-SMA genetic mutations predisposing the aortic wall to dilations and tearing, when challenged with increased hemodynamic load induced by ang II-hypertension. In addition, our finding of increased apoptotic VSM cells in aneurysmal aortas of angII-treated BSMKO mice (Figure 8B) is consistent with a pro-apoptotic effect of Bcl11b deletion in lymphoblastic T-cells by affecting p53 ^58^, a pro-apoptotic transcription factor whose uncontrolled activation has been causally linked to aortic aneurysms and dissection ^59^. Moreover, lack of VSM Bcl11b may interfere with VSM SirT1 function, a lysine deacetylase known to interact directly with Bcl11b ^50^ and p53 ^60^, and that we previously showed being indispensable in the aortic wall to prevent obesity-induced arterial stiffness ^11^ and angII-induced aortic dissections ^12^. Lastly, although the clinical relation between arterial stiffness and aortic aneurysms remains controversial, recent clinical evidence demonstrated a direct correlation between PWV and aneurysm development or progression in individuals genetically predisposed to aortic aneurysm ^61,62^. Therefore, increased aortic stiffness may have contributed to aortic mechanical failure (i.e., aortic dilatations and ruptures) in BSMKO mice when challenged with an increased hemodynamic load (angII-induced hypertension).

In conclusion, our study has uncovered a novel and crucial role for VSM Bcl11b in aortic structural and functional integrity and strongly supports Bcl11b as a potential therapeutic target against arterial stiffness and aortic aneurysms. Further studies in human populations are warranted to establish whether the genotype at the 3’-*BCL11B* locus may be used as diagnostic biomarker to identify individuals at increased risk of developing arterial stiffness and other vascular diseases.

## FUNDING SOURCES

This work was partially supported by NHLBI grant HL105287, NIA grants AG053274 and AG050599, NHLBI’s Framingham Heart Study Contracts No. N01-HC-25195 and HHSN268201500001I and grants HL070100, HL080124, HL107385 and HL126136, CTSI pilot grant 1UL1TR001430 and by the Evans Center for Interdisciplinary Biomedical Research Arterial Stiffness ARC at Boston University (http://www.bumc.bu.edu/evanscenteribr/).

## AUTHORS CONTRIBUTIONS

JACV, PE, CN, KS performed experiments and reviewed the manuscript; DA provided mouse strains used in the study; RAC, GM and KGM provided critical comments to the study and the manuscript; FS contributed to the study design, coordinated the study, designed and performed experiments, analyzed and interpreted the data and wrote the manuscript.

## DISCLOSURES

GM is the owner of Cardiovascular Engineering Inc., a manufacturer of biomedical devices for arterial stiffness measurements.

